# Chemical Environment of the Confinement Governs Thermal Stability of the Folded Telomere G-Quadruplex

**DOI:** 10.1101/2025.05.08.652821

**Authors:** Trideep Majumdar, Asim Bisoi, Prashant Chandra Singh

## Abstract

In this study, the effect of the chemical nature of the confinement on the folding and thermal stability of the telomere G-quadruplex (G4) has been investigated by studying the folding pattern of different telomere DNA sequences with varying numbers and arrangements of thymine loop nucleobases in the presence of anionic and cationic nanosized water pools. The findings suggest that both anionic and cationic water pools fold the telomere sequences into G4 of the same topology. However, the thermal stability of the folded G4 in the cationic water pool is significantly lower than that of the anionic case. The overall data indicate that the topology of the folded G4 is insensitive to the nature of the confinement, however, the thermal stability of the folded telomeric G4 depends significantly on the chemical nature of the confinement. It is plausible that the interfacial water inside the cationic water pools has a different orientation and hydrogen bonding than the case of anionic water pools, which may cause the different thermal stability of the G4 on these two water pools. These findings may be important in understanding the folding and stability of telomere G4 inside the confined cellular system.

## Introduction

The cellular system inherently possesses a confined space, which affects the folding and unfolding processes of proteins and nucleic acids, such as G-quadruplexes (G4) and i-motifs.^1-6^ Earlier experiments by Hartl and co-workers have shown that protein folding is facilitated by confinement^7-8^, a phenomenon described as the confined volume effect^9-10^, in which the entropy of the unfolded state decreases compared to the folded state.^11-13^ Recently, Mao and coworkers have demonstrated that the stability of duplex DNA decreases under the confinement effect. ^14^ In contrast, confinement induces the folding of G4 structures and stabilizes them compared to open space.^14^

The confined space plays a crucial role in telomere DNA elongation during the synthesis of 5′-GGGTTA repeat units.^6^ Therefore, it is essential to understand the folding of telomere sequences into G4 structures within such a confined environment. Mao and coworkers have demonstrated the telomere G4 folding in the non-interacting confinement environment.^6^ In this study, interactions between the confined molecules and the telomere were eliminated to exclusively understand the confinement effect on telomere G4 folding. However, in reality, telomere DNA may interact with the confined system, potentially affecting the folding process differently than in a non-interacting confined environment. Thus, it is important to investigate the effect of the chemical nature of confinement on the folding of telomere sequences into G4 structures.

Reverse micelle (RM) has been used as a model system to mimic the confinement of the cell size water droplets.^15-18^ It has been shown that RMs formed by the anionic surfactant Aerosol OT (AOT, sodium bis(2-ethylhexyl) sulfosuccinate, Figure 1a) induce the folding of a specific telomere (tel21) sequence into G4 structures.^15, 18^ Therefore, in this study, we investigated the folding and thermal stability of telomere sequences by varying the number and arrangement of thymine loop nucleobases (Figure 1b) in anionic AOT-based and cationic cetyltrimethylammonium bromide (CTAB, Figure1a)-based RMs with different water contents to understand the role of the chemical nature of confinement in the folding and thermal stability of telomere DNA sequences into G4 structures. The earlier study on the folding of the telomere sequences in the presence of the ions indicates that the folding and thermal stability of the telomere G4 is sensitive to the number and arrangement of the thymine loop nucleobases.^19-25^ Hence, varying the number and the arrangement of the thymine loop nucleobases of the telomere in the AOT and CTAB RMs will provide the generalized finding about the role of the chemical nature of the confinement on the folding and thermal stability of the telomere DNA sequences.

**Figure 1.**
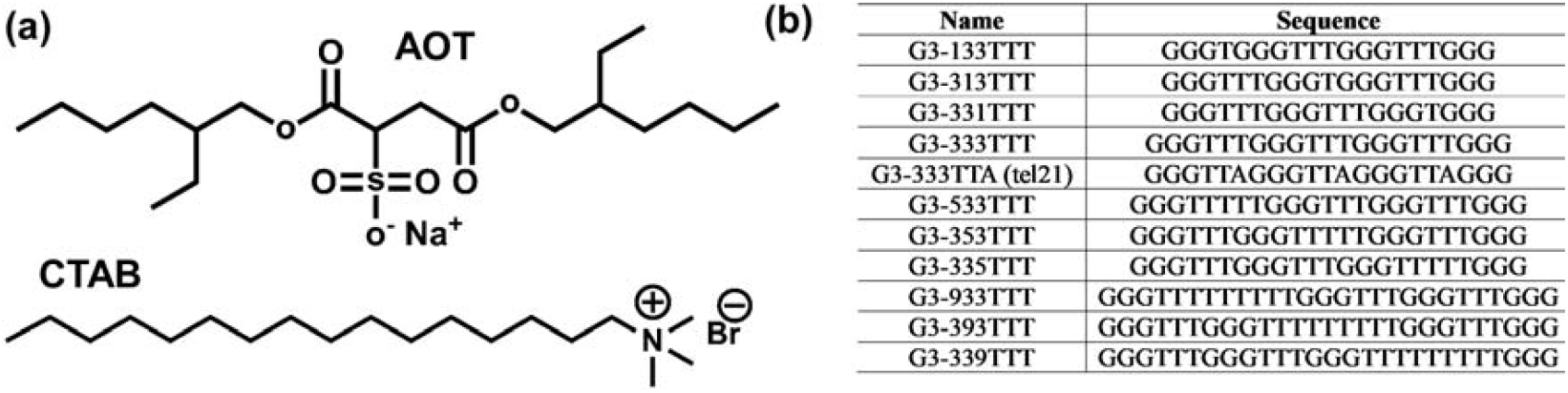
(a) The structure of the anionic surfactant AOT and cationic surfactant CTAB used for the preparation of RMs. (b) The DNA sequences used in this study and their abbreviated name.

The findings indicate that the topology of confinement-induced folded G4 structure of telomere DNA depends on the number of thymine loop nucleobases and is independent of the arrangement. Both AOT and CTAB RM fold the telomere DNA sequences into the same topology of G4. In contrast, the thermal stability of the folded G4 for telomere DNA sequences is significantly less in the case of the CTAB-RM than AOT-RM. These results suggest that while the topology of the folded G4 structure is insensitive to the nature of confinement, the thermal stability of telomeric G4 structures is strongly influenced by the chemical nature of the confinement.

## Results and Discussion

The fluorescence emission data of the FAM (donor) and TAMRA (acceptor) attached dye with G3-133TTT DNA in Tris HCl-buffer (further denoted as buffer) and the presence of AOT-RM (Figure 2a) with different water content per surfactant molecule (Wo) indicate that the fluorescence intensity of the donor of G3-133TTT decreases in the presence of AOT-RM than the buffer. The calculated FRET data suggests that the presence of AOT-RM increases the FRET efficiency between FAM-TAMRA than the buffer (Figure 2b), thereby decreasing the distance between the donor and acceptor attached to G3-133TTT compared to the buffer (Figure S1). The FRET data indicate that the G3-133TTT gets folded in the presence of AOT-RM. The steady-state FRET data of all other DNA sequences with the variation in the number of thymine loop nucleobases shows the same trend as the G3-133TTT, indicating that AOT-RM induces the folding of all the DNA sequences with the variation of the thymine in the loop region (Figure S1). To corroborate the steady state FRET data, the lifetimes of the donor (FAM) and acceptor (TAMRA) attached with DNA sequences in buffer and AOT-RM were measured. The lifetime of the donor attached with DNA sequences decreases, while the acceptor lifetime increases in the presence of AOT-RM (Table S1), confirming FRET between the donor and acceptor due to the folding of these DNA sequences in the presence of AOT-RM (Figures 2c-d, S2). Interestingly, the FRET efficiency of all the DNA sequences is maximum for 5Wo and decreases with the increase of the water content with minimum FRET for 40Wo, regardless of the number of thymine loop nucleobases (Figures 2 a-d). The data indicate that the folding extent of all the DNA sequences in the presence of AOT-RM depends on the number of water molecules in the RM, in line with the earlier finding for the tel21 DNA sequence.^18^

**Figure 2.**
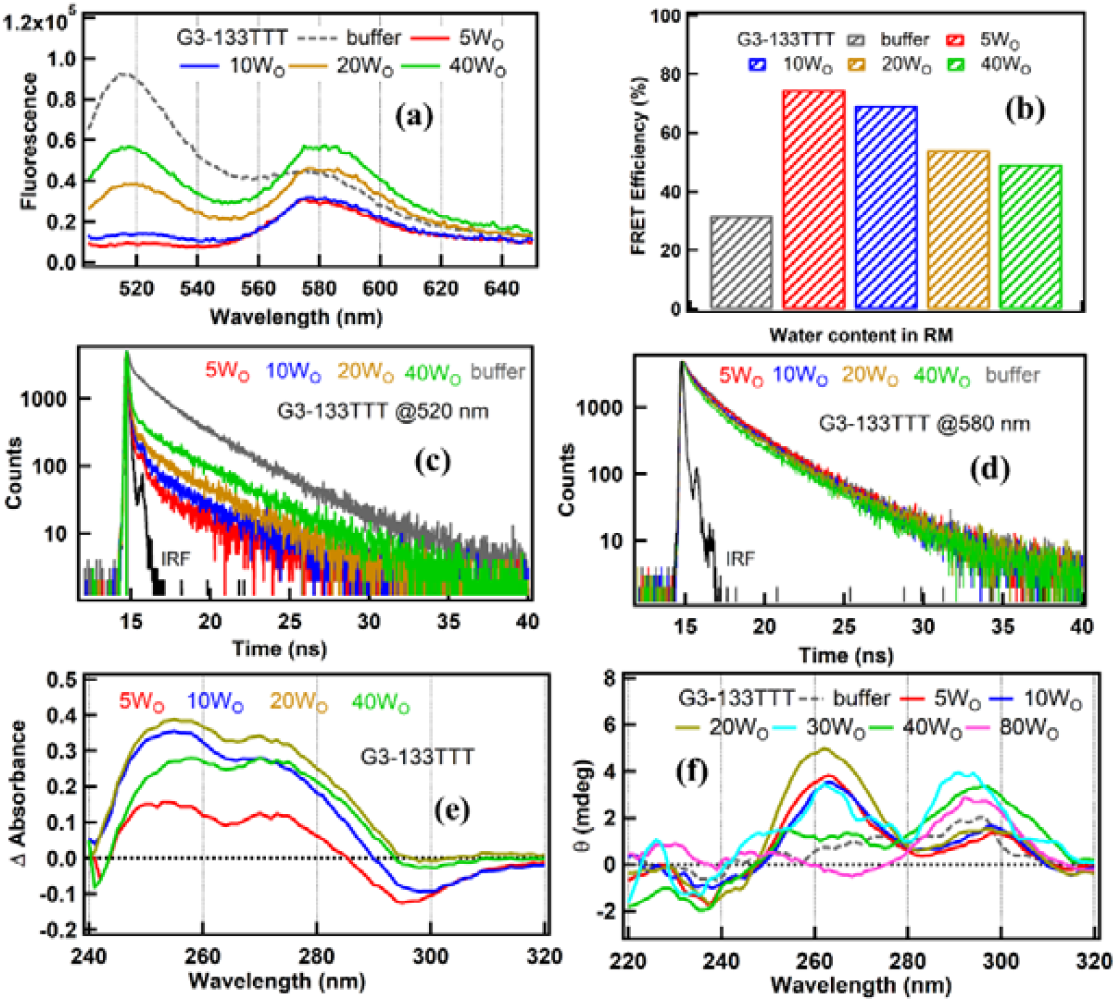
(a) The fluorescence spectra of the FAM (donor, 520 nm)-TAMRA (acceptor, 580 nm) attached with G3-133TTT DNA (25 nM) in buffer and the AOT-RMs having different water content. The fluorescence spectra have been obtained by exciting at the absorption peak maxima of FAM at 492 nm. (b) The FRET efficiency of the FAM-TAMRA attached G3-133TTT DNA (25 nM) in buffer and the AOT-RMs having different water content. (c) The lifetime decay traces of the FAM of the FAM-TAMRA attached G3-133TTT DNA (25 nM) in buffer and the AOT-RMs having different water content obtained from the excitation of the FAM at 450 nm. (d) The lifetime decay traces of the TAMRA attached G3-133TTT DNA (25 nM) sequence in buffer and the AOT-RMs having different water content obtained from the excitation of the FAM at 450 nm. (e) The isothermal difference spectra (IDS) of the G3-133TTT (5 µM) in the presence of the AOT-RM of different water content. The IDS spectra were obtained by subtracting the absorbance spectra of the G3-133TTT in buffer and the presence of the AOT-RM of different water content. (f) The CD spectra of G3-133TTT (5 µM) in buffer and AOT-RM of the different water content. All the measurements were performed at 22±2□.

The isothermal difference spectra (IDS) of the G3-133TTT and other DNA sequences were obtained by subtracting the absorbance spectra of the G3-133TTT in buffer from those in the presence of RM to understand the nature of the folded DNA in AOT-RM (Figure 2e). The IDS spectra of the G3-133TTT show the negative peak at 295 nm, suggesting the possibility of folding the G3-133TTT into G4^26-27^. A similar trend of the IDS spectra has been observed for all other sequences in AOT-RM (Figure S3), indicating the possibility of folding these sequences into G4 in the presence of AOT-RM.

Further, the CD spectra of all these DNA sequences in buffer and the AOT-RM of different water content have been performed to understand the topology of the folded DNA structure in the presence of RM. The CD spectra of G3-133TTT (Figure 2f) in AOT-RM of 5-20 Wo depict the positive ellipticity peak at 260 nm, along with a negative ellipticity peak around 240 nm. A small 290 nm positive ellipticity peak is also observed; however, its ellipticity is significantly less than at 260 nm. The CD feature of G3-133TTT in the presence of RM of 5-20 Wo is similar to its CD spectrum in KCl, known to form a parallel structure (Figure S4).^19^ With a further increase in the water content of the RM, the spectral feature of the CD of G3-133TTT changes as the positive ellipticity of the 260 nm decreases and 290 nm increases monotonically. The ellipticity of the peak at 290 nm is higher than 260 nm with the addition of RM of 30 Wo, similar to the hybrid type G4 (Figure S4). The CD spectral feature of G3-1333TTT in RM of 40Wo shows mainly a 290 nm peak with an almost zero ellipticity peak at 260 nm. With the further increase of water content in RM (RM=80Wo), the CD spectrum of G3-133TTT depicts the positive ellipticity peak at 290 nm and negative ellipticity feature at 260 nm, similar to the case of the CD spectra of G3-133TTT in NaCl (Figure S4) known to form the antiparallel structure.^28^ The CD data indicate that G3-133TTT in RM shows the conformation change in the folded G4 depending on the water content in RM as it forms the parallel G4 for 5-20Wo and then folds into hybrid and antiparallel topology with further increase of the water content in RM. The spectral feature of the CD data of the G3-313TTT and G3-331TTT is the same as the G3-133TTT (Figure S5) in the AOT-RM, indicating that the arrangement of the thymine nucleobases in the loop region does not affect the confirmation of the folded G4.

In contrast to G3-133TTT, the CD spectra of G3-333TTT, and G3-533TTT in AOT-RMs of all the Wo exhibit the positive ellipticity peaks at 290 nm along with a negative ellipticity peak around 265 nm (Figure 3a) which is closer to the CD data of same DNA sequences in the presence of NaCl (Figure S4), known to form the antiparallel G4 structure.^28^ The CD data of G3-333TTA (tel21) in RMs suggests that it folds into antiparallel topology for all the Wo in agreement with earlier studies,^18^ similar to the G3-333TTT and G3-533TTT sequences (Figure 3a). The CD data of G3-933TTT in RMs for all Wo show the positive ellipticity feature at 290 and 260 nm, along with the negative ellipticity peak at 240 nm (Figure 3a) indicating the formation of hybrid G4 as observed in the case of both KCl and NaCl salts for the same sequence (Figure S4).^29^ The spectral feature of the CD data of all these sequences in RM does not change significantly on altering the arrangement of the thymine nucleobase in the loop, as observed in the case of G3-133TTT (Figure S5). The confirmation changes of the G4 in RM depending on the water content is specific only for the G3-1333TTT series of DNA sequence whereas, in all other cases, the topology of the folded G4 is the same for all the Wo. Nevertheless, in general, the CD data for all DNA sequences with varying thymine loop nucleobases indicate that the topology of the folded G4 structure is sensitive to the number of thymine nucleobases in the loop region but is insensitive to variations in their arrangement.

**Figure 3.**
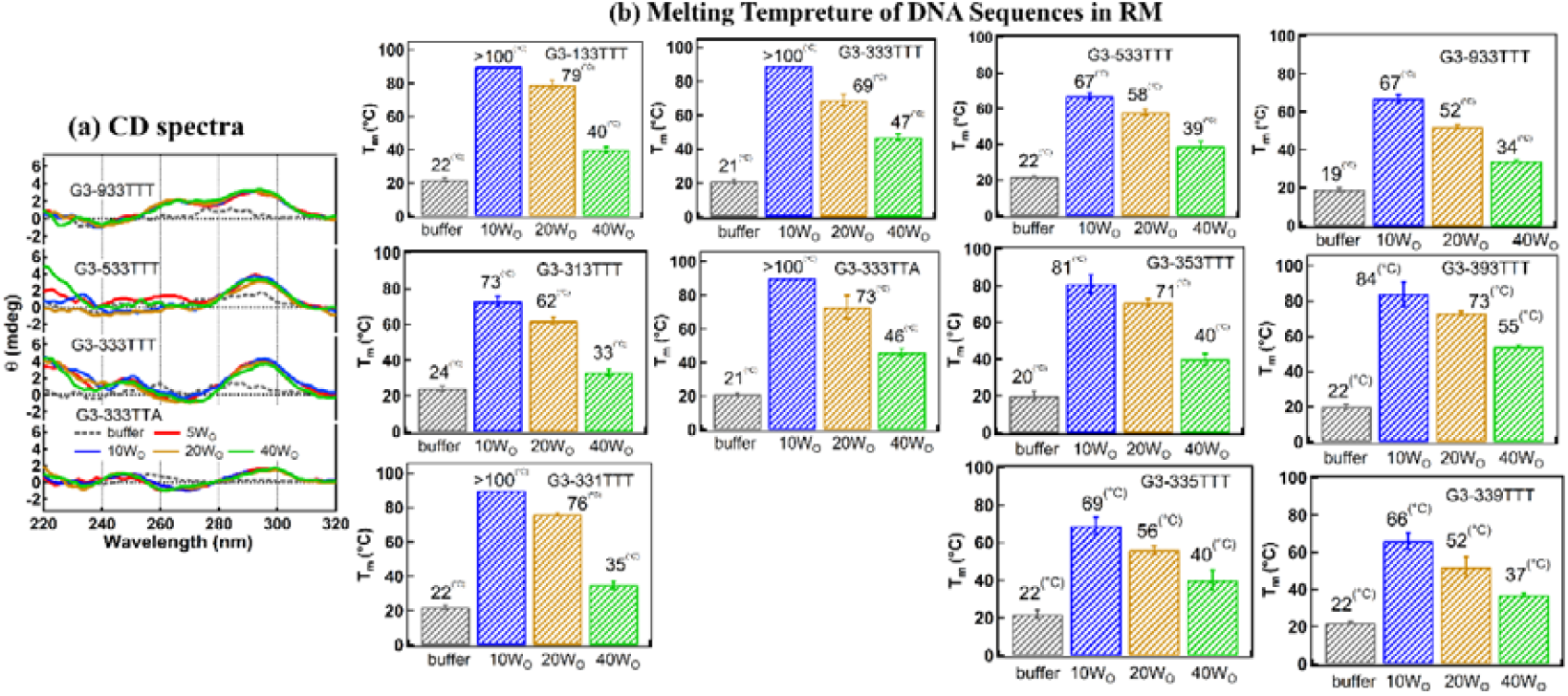
(a) The CD spectra of the different DNA sequences (5µM) in buffer and the presence of AOT-RM with different water content. (b) The melting temperature of the DNA sequences (5µM) in buffer and the presence of the AOT-RM of different water content.

To understand the effect of the AOT-RM on the thermal stability of the folded G4 structure of G3-133TTT, temperature-dependent CD experiments have been performed. The change in the ellipticity at the characteristic G4 peak (290 nm in the case of hybrid and antiparallel G4, whereas 260 nm in the case of parallel G4) ^30-32, 20, 33-41^has been plotted against temperature to estimate the melting temperature (T_m_) of these DNA sequences in the presence of AOT-RM with different water content (Figure S6). The T_m_ of DNA sequences in the RM=5Wo is more than 100□, as the folded G4 structure could not melt completely within the measurement range of the experiment up to 100□. The T_m_ of all DNA sequences in the case of 80Wo could not be measured as the RM=80Wo itself gets broken at higher temperatures (Figure S7). Nevertheless, in general, the T_m_ of all the DNA sequences decreases with the increase of the water content in the AOT-RM as the T_m_ is significantly less for the 40Wo than in the case of 10Wo (Figure 3b) which is in agreement with the FRET data indicating that the folding efficiency of DNA sequences decreases with the increasing water content in RM. Interestingly, while the topology of the folded G4 structure is insensitive to the arrangement of nucleobases, the T_m_ of the folded G4 structure depends on the arrangement of loop nucleobases within the DNA sequences. For example, the thermal stability of the folded G4 for the DNA sequences with the short loop (G3-133TTT) is higher when the small loop is present at the terminal (G3-133TTT and G3-331TTT) rather than at the central position (G3-313T). In contrast, the thermal stability of the folded G4 of the DNA sequences with longer loop length (G3-533TTT and G3-933TTT) is maximum when the loop nucleobases are present in the central position (G3-353TTT, G3-933TTT) rather than the terminal place (Figure 3b). This study suggests that apart from the number and arrangement of thymine loop nucleobases, the amount of water inside RM (which corresponds to the RM size) plays a significant role in the thermal stability of the folded G4 structure in DNA sequences with different thymine loop configurations. Notably, the thermal stability of the folded G4 structure in AOT-RM with water content below 40 Wo is higher than in the presence of either Na□ or K□ ions, indicating that AOT-RM with fewer water molecules provides greater stability to the G4 structure than these known ion cases.^19, 28^ Overall, these findings indicate that the topology of the folded G4 in the AOT-RM depends on the number of loop nucleobases rather than its arrangement whereas the thermal stability of the folded G4 depends on the number of water molecules in the RM, number and arrangement of the thymine loop nucleobases in the DNA sequences. The dependency of the melting temperature of G4 in AOT-RM on the arrangement of the thymine nucleobases suggests that the local interaction between the nucleobases of the same topology of G4 may be different due to the possibility of the different polymorphic structures of the same topology of G4.^42^

Next, the CD study of the DNA sequences mentioned above in the presence of the CTAB-RM has been performed (Figure 4) and compared with AOT to understand the role of the chemical nature of the headgroup (AOT is negatively charged, and CTAB is positively charged) in the folding of these sequences into G4. The G3-133TTT depicts the parallel topology of G4 in CTAB-RM, which changes towards the antiparallel type in the presence of the higher water content of CTAB-RM (Figure 4a). The G3-333TTT and G3-533TTT form an antiparallel topology of G4, whereas G3-933TTT gets folded into the hybrid topology in the presence of CTAB-RM irrespective of the water content (Figures 4 b-d). The topology of the G4 in the CTAB-RM does not depend on the arrangement of the thymine loop nucleobases as observed in the case of AOT-RM. The topology of the G4 of different DNA sequences in the presence of CTAB-RM is similar to that in AOT-RM, indicating that the chemical nature of the confinement does not significantly influence the topology of the folded G4. The dynamic light scattering (DLS) data indicate the average hydrodynamic diameter of the DNA in the presence of the RM (∼3-10 nm) decreases compared to the buffer case (∼400-600 nm), reassuring the folding of the DNA sequences in the presence of the RM (Figures 5a-b, S8, and S9). Interestingly, the hydrodynamic size of the DNA in RMs is approximately the same as that of bare RMs without DNA (Figures 5a-b, S8, and S9). The DLS data indicate that the folding of these sequences mainly occurs on the periphery of the RM, suggesting that the confinement provided by the RM is the primary factor for triggering the folding of the DNA sequences into G4, irrespective of the different chemical natures of these two RMs.

**Figure 4.**
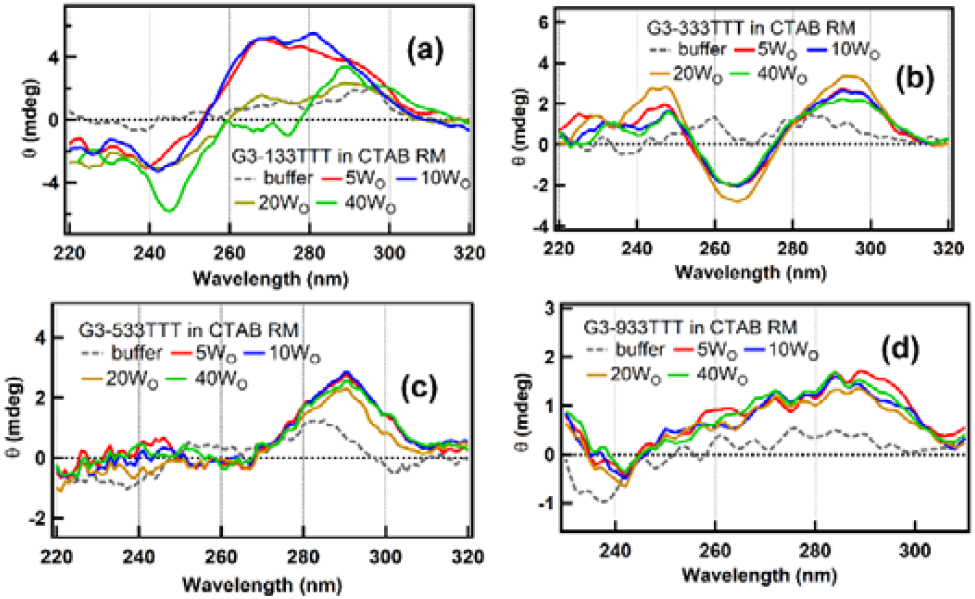
The CD spectra of G3-133TTT (a), G3-333TTT (b), G3-5333TTT, and G3-9333TTT (5 µm each) in CTAB-RM with different water content.

**Figure 5.**
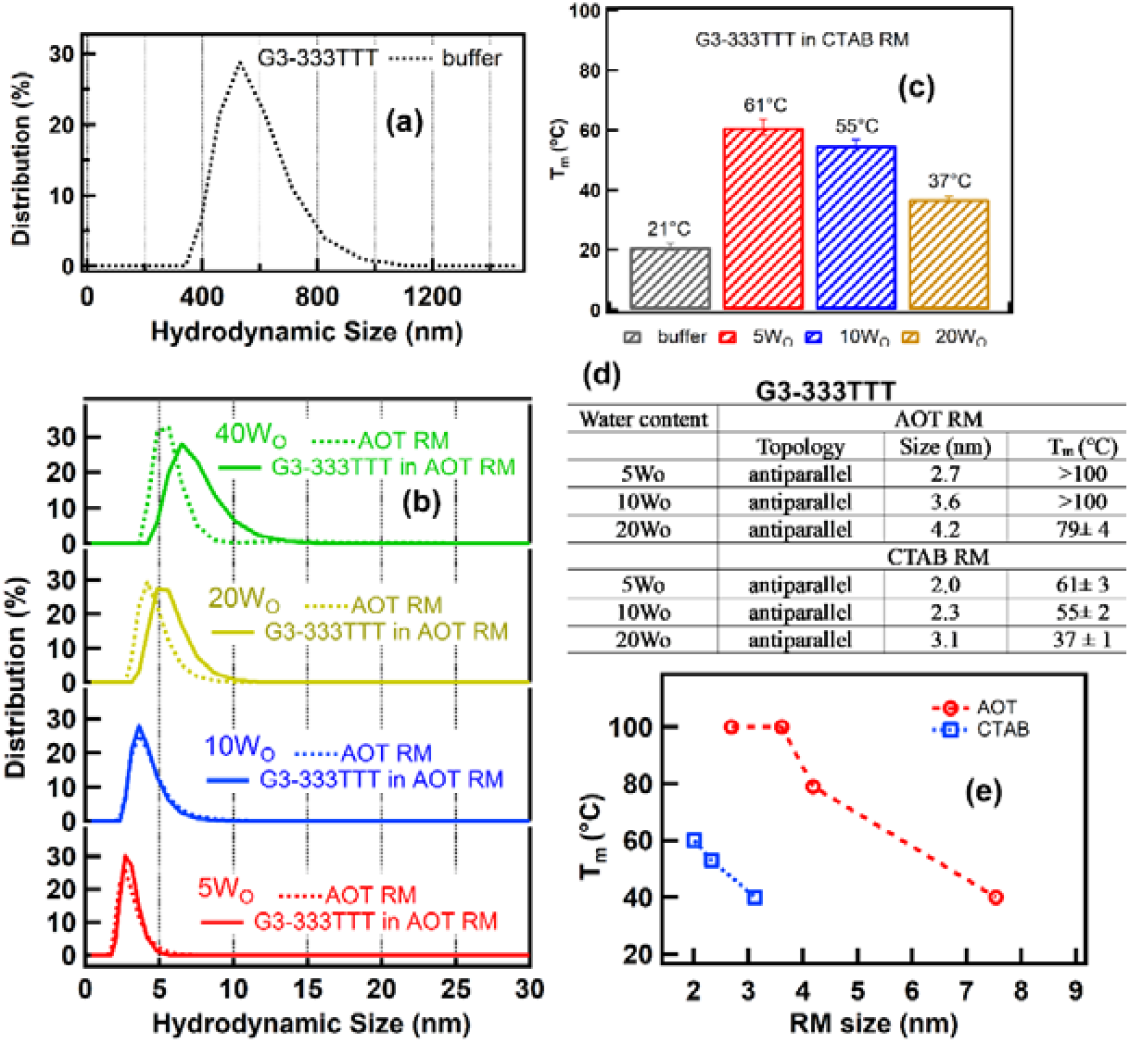
**(a)** The DLS spectra of G3-333TTT (5 µM) in buffer. **(b)** The DLS spectra of AOT-RM of the different water content in the absence and presence of G3-333TTT DNA (5 µM) sequence. (c) The melting temperature of G3-333TTT DNA sequence (5 µM) in the presence of CTAB-RM with different water contents (d) The comparison of the topology and the melting temperature of G3-333TTT in presence of AOT and CTAB RMs (e) The representation of the variation of the melting temperature of the G4 of the G3-333TTT in AOT and CTAB RMs.

Finally, the melting temperature of the DNA sequences in the CTAB-RM has been estimated to understand the stability of the telomeric G4 in the presence of cationic RM. The melting temperature of the G4 in the CTAB-RM for higher water content (Wo≥40) could not be determined accurately as it melts rapidly and produces the noisy temperature-dependent CD spectra. Nevertheless, the trend of the melting temperature data indicates that the thermal stability of G4 decreases with the increasing confinement size in both RMs, indicating that the size of the confinement is a critical player in the thermal stability of the G4 in the presence of RMs (Figures 3b and 5c-e, S10). Small-sized RMs (5–10 Wo) contain mostly interfacial water, whereas larger-sized RMs (≥20 Wo) have significant bulk water contributions. It is known that the nature of the interfacial water is distinctly different from the bulk in terms of the hydrogen bonding and the orientation.^43-44^ Moreover, the motion of the interfacial water near the surfactant head group of the RM is more restricted than the bulk water, which may be one of the few reasons for the decrease in the thermal stability of the G4 with the increase in the confinement size. Indeed, in one of the earlier studies, it was discussed that the activity of the water in the nanoconfinement decreases, promoting G4 folding in the confinement environments.^3^

Interestingly, despite the similar trend of thermal stability, the absolute value of the melting temperature of G4 in the case of the CTAB-RM is significantly less than AOT-RM (Figures 5d-e, S10, Table S2). The hydrodynamic size of the CTAB-RM for a fixed Wo is similar or even slightly less than the AOT-RM (Figure S11). Therefore, according to the size of the confinement, the thermal stability of the G4 in the CTAB-RM should be approximately the same as the AOT-RM, which contradicts the experimental findings. Hence, the trend of the thermal stability of the telomere G4 in AOT and CTAB-RMs indicates that the role of the chemical nature of the surfactant is also important in the thermal stability of the G4 apart from the confinement effect. It is known that the positively charged surfactant affects the orientation and hydrogen bonding between the interfacial water and the interaction of water with the head group differently than the negatively charged.^44^ Hence, it can be speculated that the different thermal stability of the G4 at CTAB-AOT than AOT-RM may be due to the different hydrogen bonding environment and orientation of interfacial water near the CTAB-AOT than AOT-RM which affects the water activity near these two RMs in different manner.^3^ Overall, these findings indicate that the chemical nature of the confinement plays a prominent role in the thermal stability of the G4 at the RM.

## Conclusion

The telomeric DNA sequences fold into the same topology of G4 in the presence of both cationic and anionic RMs. The topology of the folded G4 depends on the number of the thymine loop nucleobases, whereas the role of the arrangement of the thymine loop nucleobases is not critical. The thermal stability of a folded G4 DNA sequence decreases with the size of both RMs. The thermal stability of the folded G4 also varies with the number and arrangement of the thymine nucleobases in telomeric G4. Interestingly, the thermal stability of the folded telomeric G4 in the presence of cationic RM is significantly less than the anionic RM, indicating that the chemical environment of the confinement plays a critical role in the thermal stability of the G4. This study suggests that the topology of the folded telomeric G4 by the confinement is not dependent much on its chemical environment, however, the thermal stability of the folded G4 depends significantly on the chemical environment of the confinement.

