## Supplementary material for "Chemical Environment of the Confinement Governs Thermal Stability of the Folded Telomere G-Quadruplex": SI

Indian Association for the Cultivation of Science

Jadavpur, Kolkata, India 700032

*

**Materials and Methods**

All the telomeric DNA sequences used in this study are high-pressure liquid chromatography grade pure and procured from Sigma Aldrich. AOT (>98% purity) and CTAB (>98% purity) were purchased from TCI Chemicals. HPLC-graded isooctane, chloroform, and hexane were obtained from Sigma Aldrich. The RMs were prepared by dissolving the AOT/CTAB in isooctane and chloroform-hexane (3:2 v/v) mixture followed by the addition of the required amount of 10 mM Tris-HCl buffer (pH=7.4) to achieve a particular Wo = [H_2_O] / [Surfactant]. All the DNA samples were dissolved in 10 mM Tris-HCl buffer (pH=7.4) and directly injected to AOT or CTAB solutions as per requirement.

The absorption spectra of each DNA sequence (~5 µM) in the buffer and the presence of the RMs were measured at room temperature using the UV-Vis spectrometer (Evolution 201). The isothermal difference spectra of the DNA sequences were obtained by subtracting the absorbance spectrum of the particular DNA in the presence of RM from those in buffer to understand the nature of the folded DNA in RM.

The fluorescein amidites (FAM) attached as donor at the 5’ terminal and carboxy tetramethyl rhodamine (TAMRA) as acceptor attached with the 3’ terminal of DNA sequences has been used to perform the fluorescence resonance energy transfer (FRET) experiment. The fluorescence emission of the FAM (~520 nm) and TAMRA (~580 nm) attached with DNA (~25 nM) has been measured using the excitation maxima of FMA at 492 nm in buffer and RMs using the fluorometer (fluorolog3). The FRET efficiency (E) and distance were calculated using equations 1 and 2, respectively.

E = F_A_/(F_D_ + F_A_) ………..(1)

E = 1 / [1 + (r / R_0_)^6^] ………..(2)

F_D_ and F_A_ are the fluorescence intensities of the donor and acceptor. R_0_ is the Forster distance (55 Å for FAM-TAMRA pair) where 50% energy transfer takes place, whereas r is the distance between donor and acceptor in the experimental condition.

The fluorescence intensity decay curves of the FAM-TAMRA attached DNA sequences (25 nM) were measured using time-correlated single-photon counting (TCSPC) setup. A picosecond diode laser of 450 nm was used as an excitation source for exciting the FAM (donor) dye attached with the DNA sequences. The emission was collected at 520 nm (emission maxima of FAM) and 580 nm (emission maxima of TAMRA) at the magic angle polarization and detected using a multiple channel photomultiplier. The typical full width half maximum of the system was measured using a liquid scatterer and found to be ∼40 ps. Obtained data were fit with a single exponential I(t) = Σ B exp(−t/τ), where τ is the fluorescence lifetime and B is the pre-exponential factor for that component of the decay. The fitting curves were deconvoluted with the instrument response profile. The quality of the fitting was judged by the calculation of the reduced chi-square (χ^2^) value and the distribution of the weighted residuals among the data channels. The obtained values of χ^2^ were close to 1 and the weighted residuals were randomly distributed around 0, indicating a high goodness of fit.

The CD spectra of each telomeric DNA sequence (5 μM) in buffer and RMs were measured in the range of 230-320 nm using a JASCO (J1500) spectrometer at 22±2℃ to understand the topology of folded telomeric G4 by RMs. The thermal stabilization of the G4 of the different sequences folded in the presence of RMs (5 μM) has been assessed by measuring the melting temperature (T_m_) of each sequence in buffer and the presence of RMs. The melting temperature of the G4 was obtained by performing the temperature-dependent CD measurement of each DNA sample in the range of 10 ℃ to 90 ℃ in the interval of 3℃ and plotting the ellipticity change in the characteristic peak of G4 (290 nm for the antiparallel and hybrid topology, and 260 nm for the parallel G4) against the temperature. The ramp rate of the temperature for the CD measurements was 1℃/minute. Each measurement is an average of three different CD spectra.

The distribution of the hydrodynamic size of each telomeric DNA sequence (~5 µM) and RMs were measured at room temperature (22±2℃) using the Malvern dynamic light scattering (DLS) spectrometer. Each measurement was repeated three times to check the reproducibility of the data.

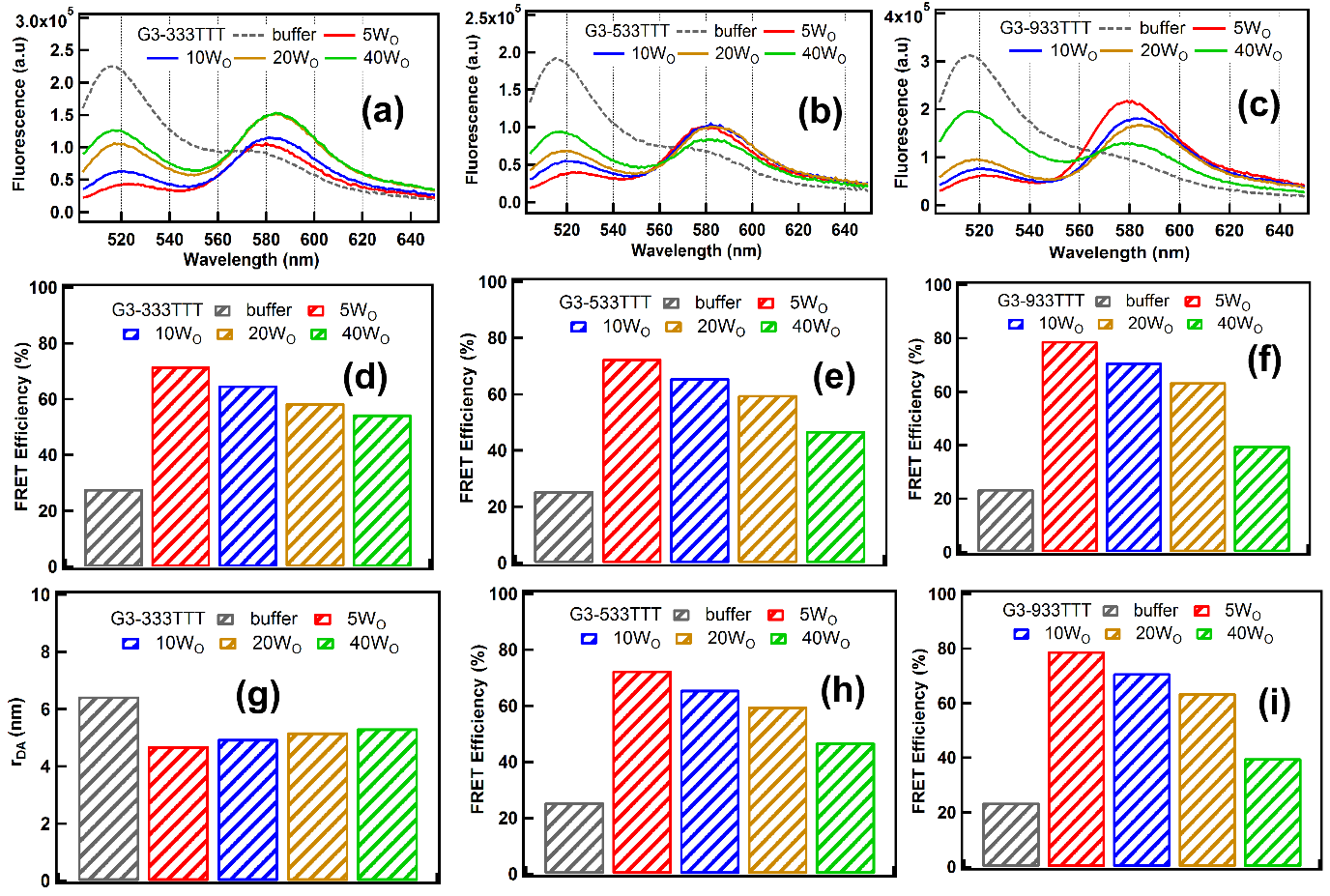

**Figure S1:** The fluorescence spectra of the FAM (donor)-TAMRA (acceptor) attached DNA sequences, G3-333TTT DNA (a), G3-533TTT (b), and G3-933TTT (c) (25 nM each) in buffer and the AOT-RMs having different water content. The fluorescence spectra have been obtained by exciting at the absorption peak maxima of FAM at 492 nm. The FRET efficiency of the FAM-TAMRA attached G3-333TTT DNA (d), G3-533TTT (e), and G3-933TTT (f) in buffer and the AOT-RMs having different water content. **(c)** The donor-acceptor distance of the FAM-TAMRA attached G3-333TTT DNA (g), G3-533TTT (h), and G3-933TTT (i) in buffer and the AOT-RMs having different water content.

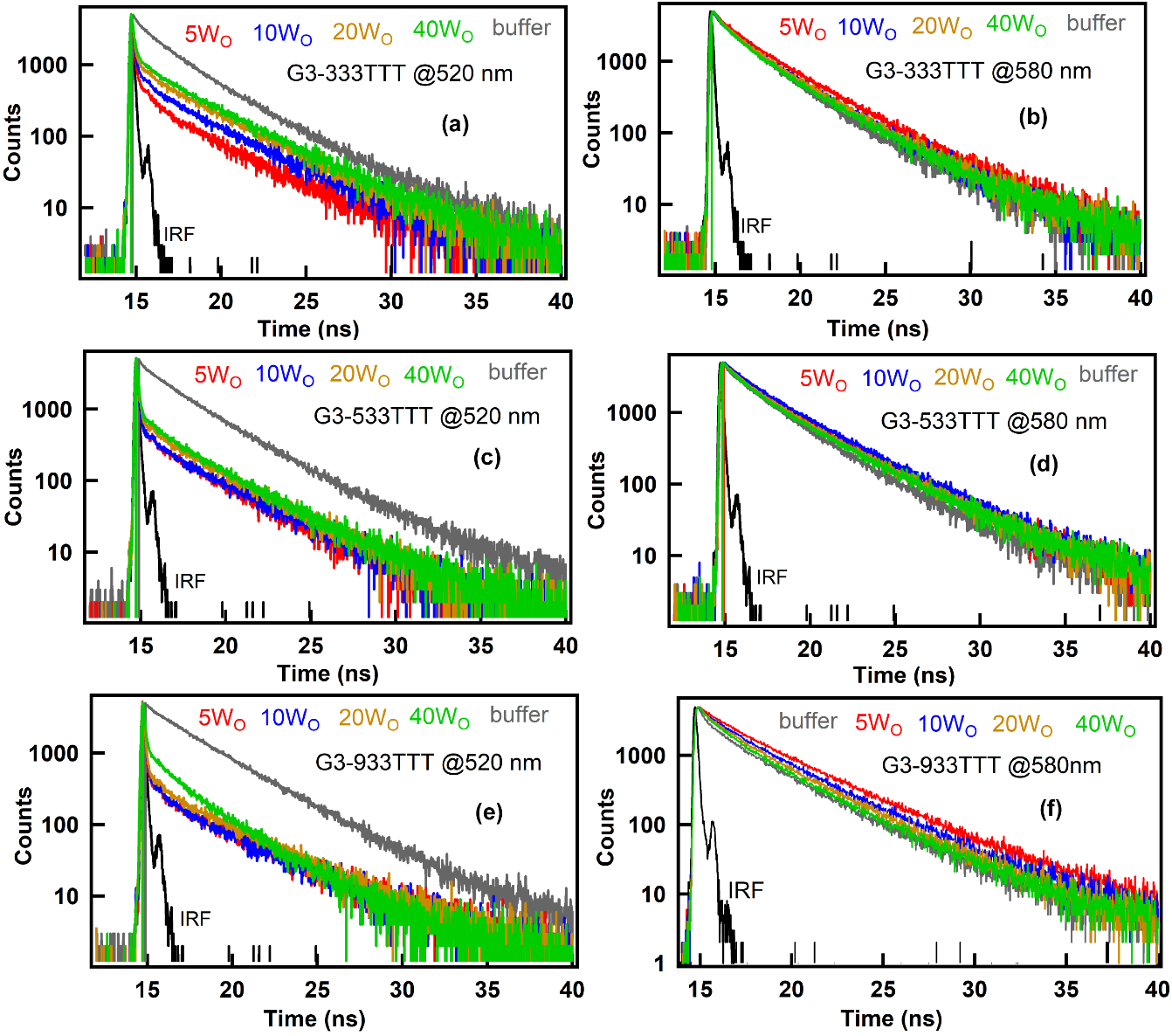

**Figure S2:** The lifetime decay traces of the FAM and TAMRA attached with G3-333TTT (a and b), G3-533TTT (c and d), G3-933TTT (e and f) DNA (25 nM) in buffer and the AOT-RMs having different water content obtained from the excitation of the absorption peak maxima of FAM.

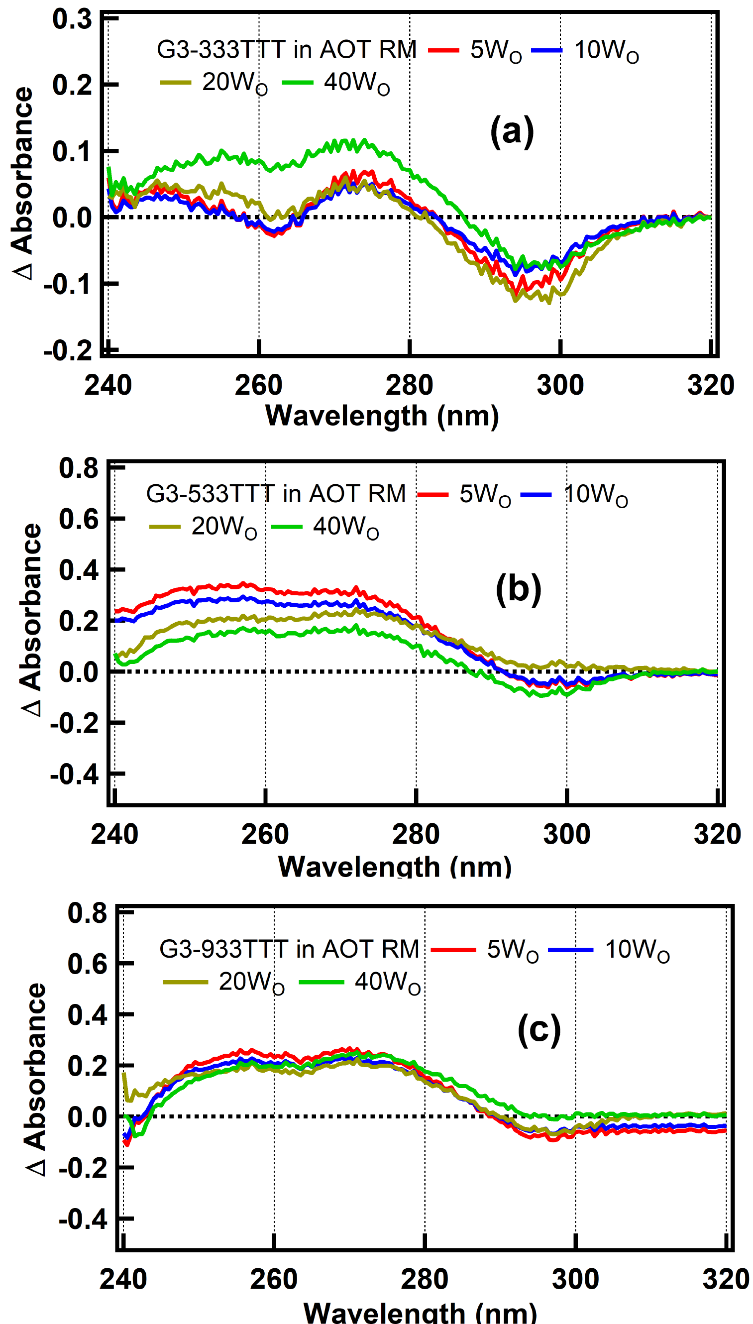

**Figure S3:** The isothermal difference spectra (IDS) of G3-333TTT (a), G3-5533TTT (b), and G3-933TTT (c) (5 µM each) in the presence of the AOT-RM with varying water content. The IDS spectra were obtained by subtracting the absorbance spectra of these DNA sequences in buffer from the absorbance spectra in the presence of the RM with different water content.

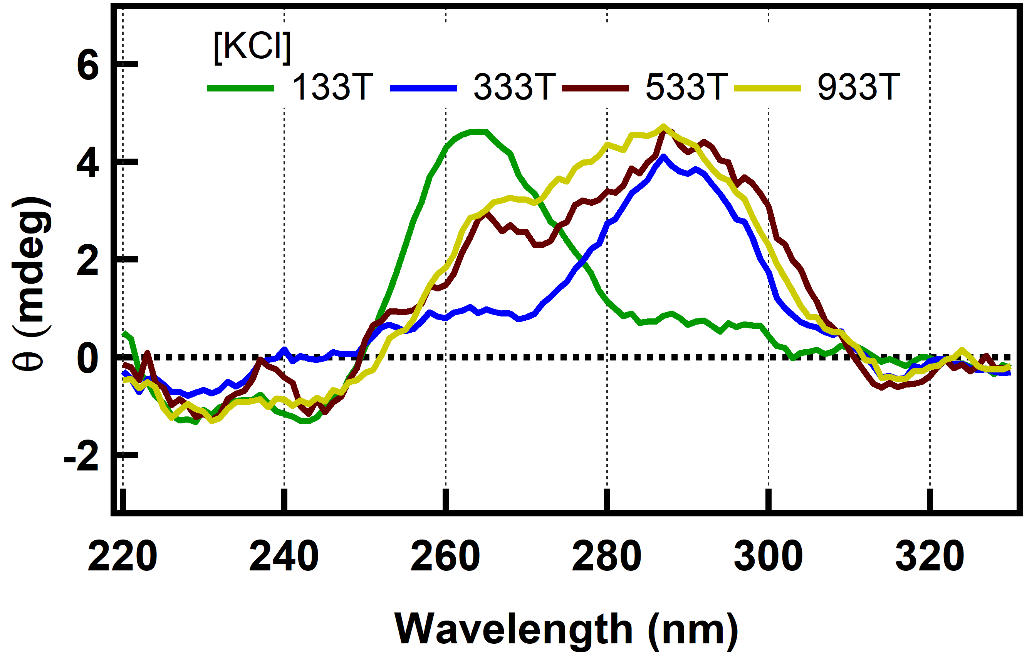

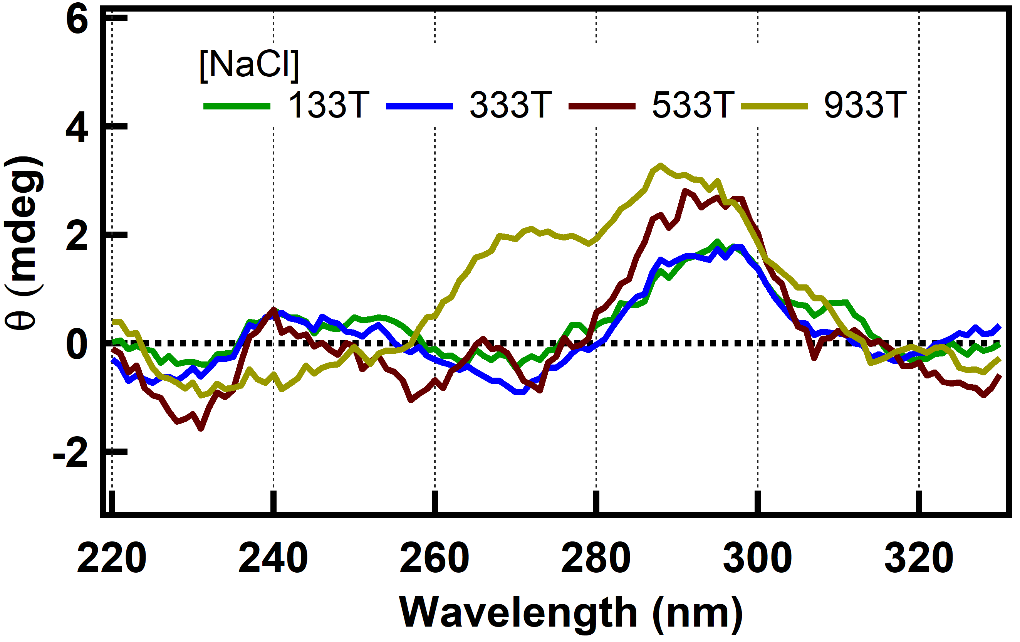

**Figure S4:** The CD spectra of different DNA sequences (5 µM each) used in this study in the presence of KCl and NaCl (100 mM each).

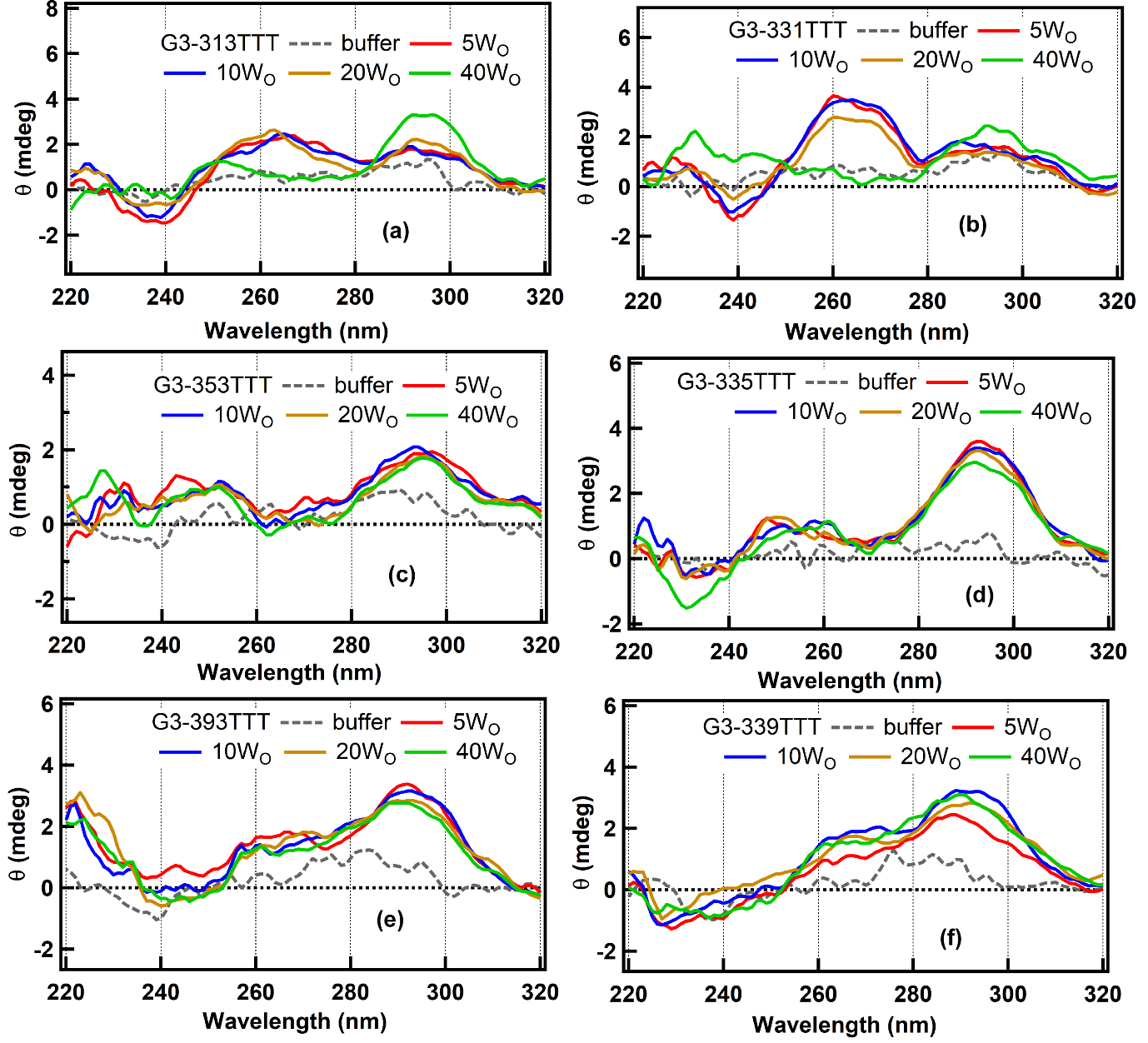

**Figure S5:** The CD spectra of G3-313TTT (a), G3-331TTT (b), G3-533TTT (c), G3-335TTT (d), G3-393TTT (e), G3-339TTT (f), (5 µM each) in buffer and RM of the different water content.
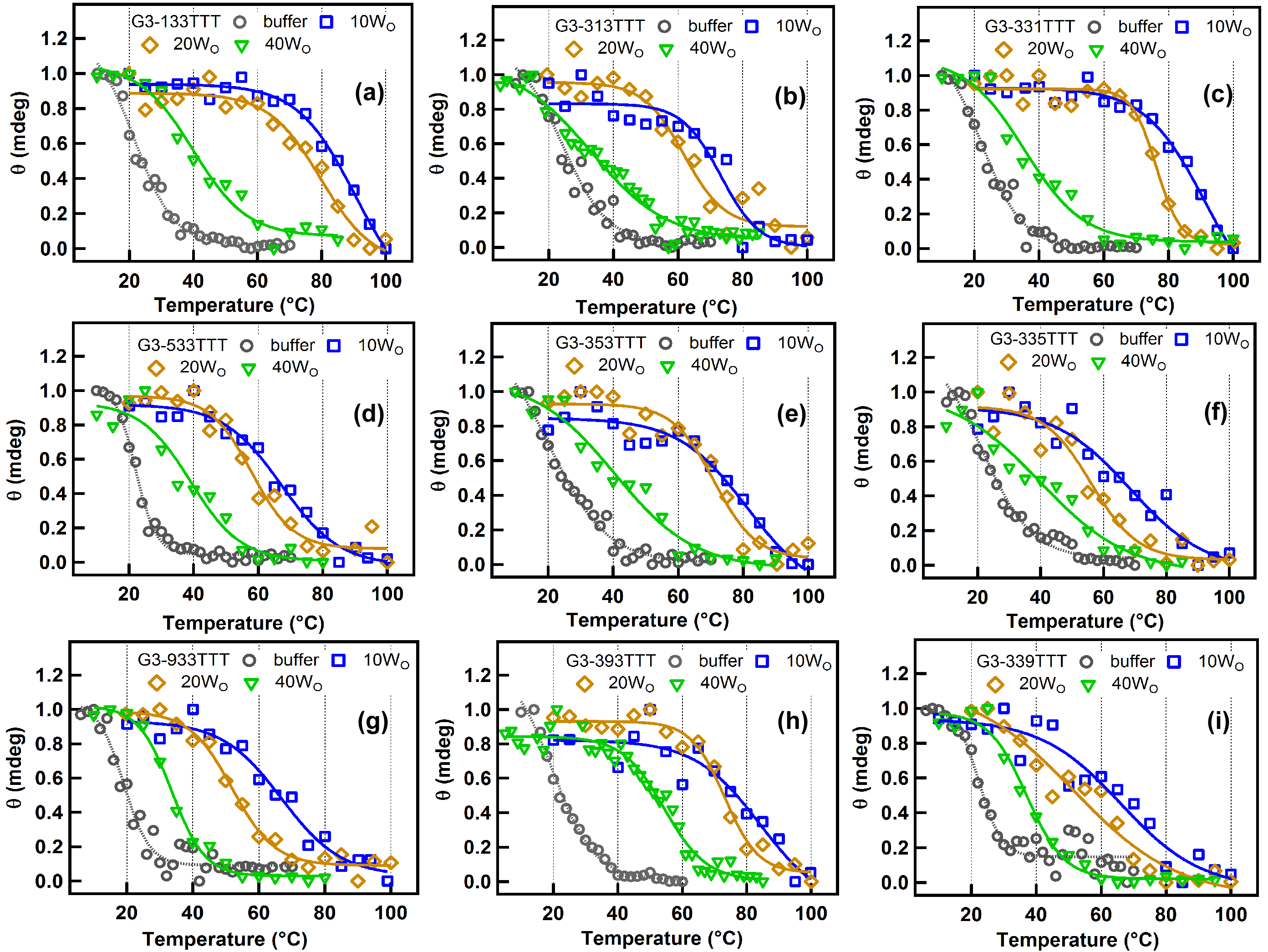

**Figure S6:** The melting curve of the different DNA sequences (5µM) in buffer and the presence of the RM of different water content.

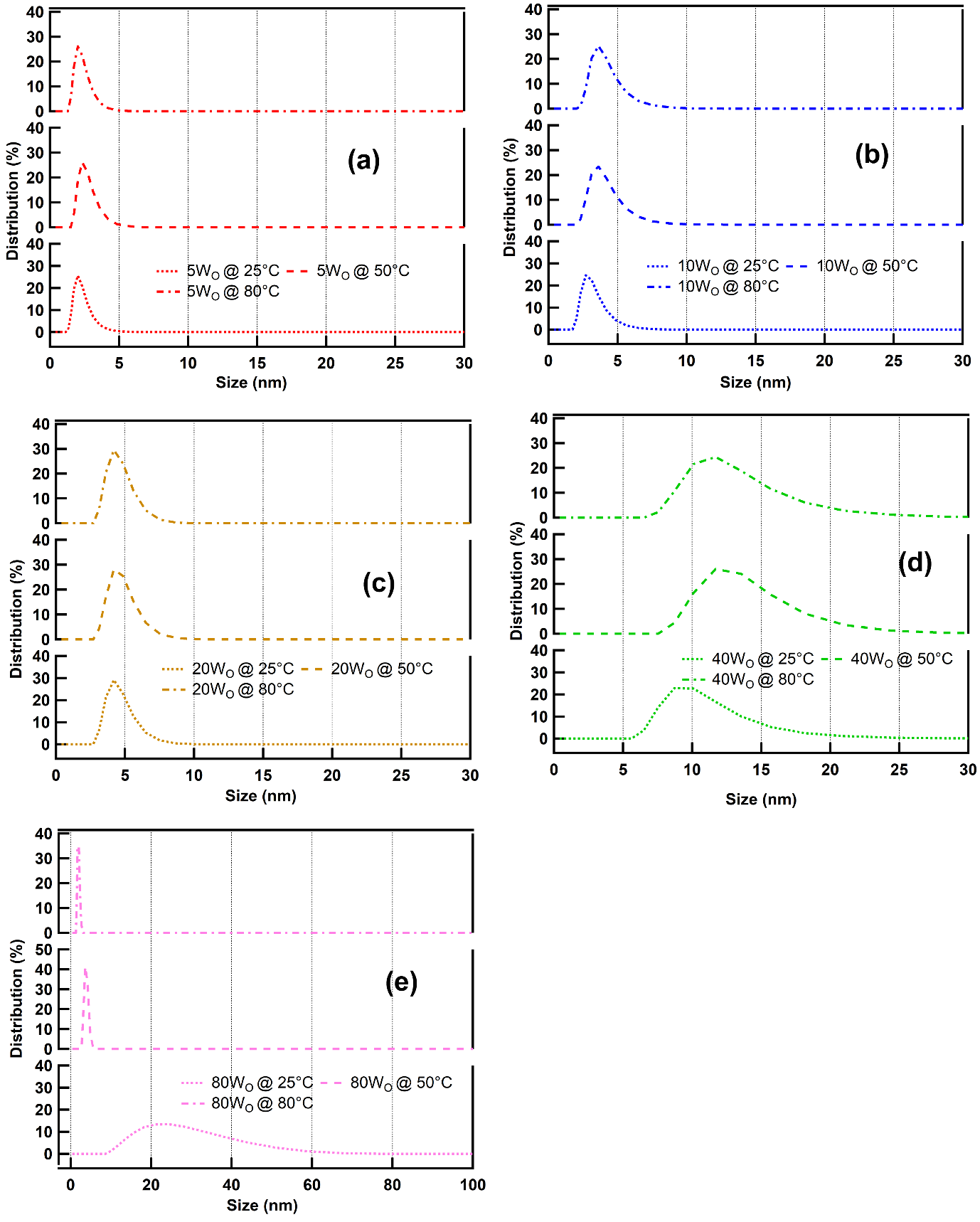

**Figure S7:** The temperature-dependent DLS spectra of the AOT RMs having (a) 5Wo, (b) 10Wo, (c) 20Wo, (d) 40Wo, (e) 80Wo, respectively.

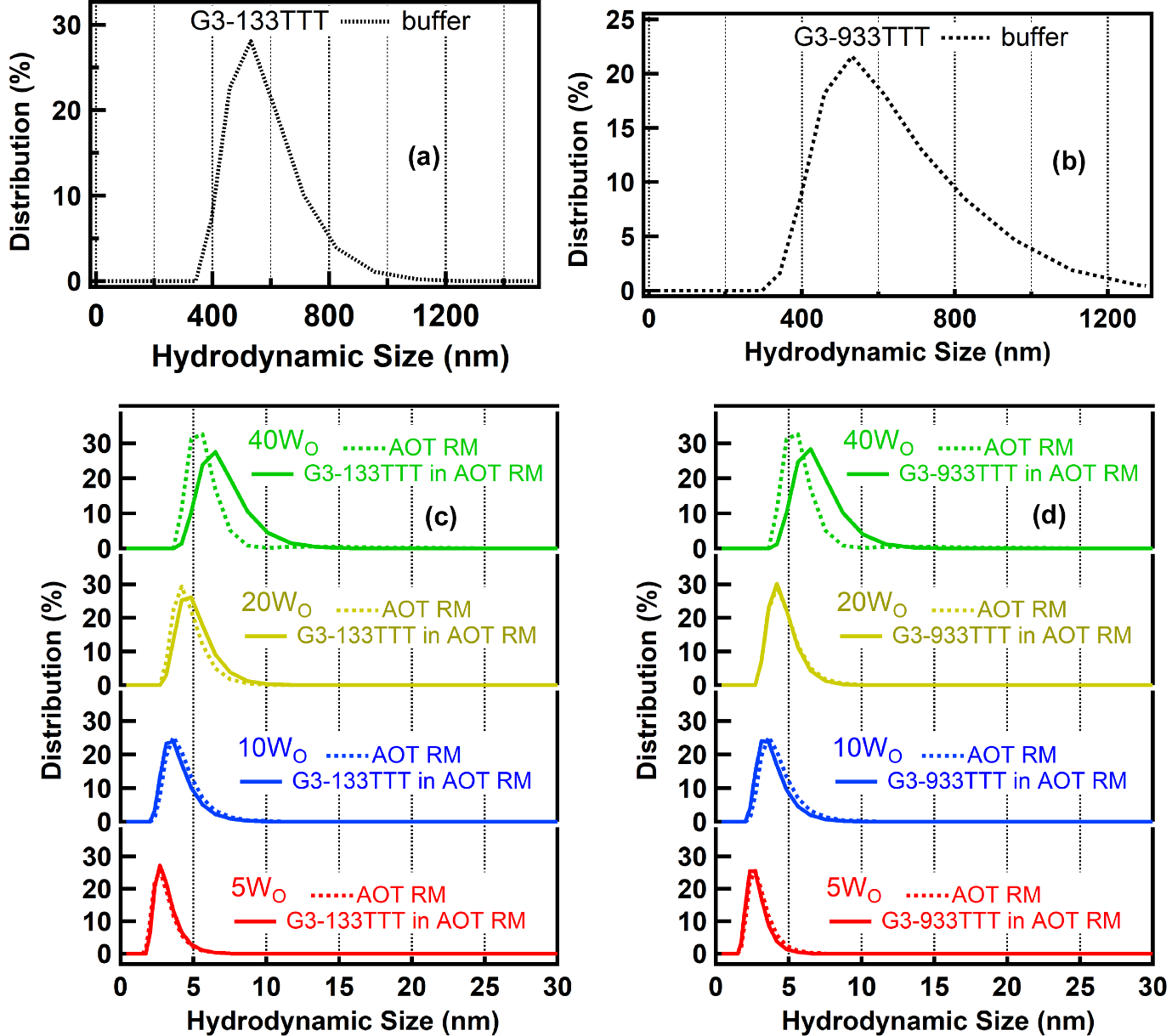

**Figure S8:** The DLS spectra of G3-133TTT (a) and G3-933TTT (b) DNA (5µM) sequences in buffer. The DLS spectra of AOT-RM without and with G3-133TTT (c) and G3-933TTT (d) DNA sequences (5µM).

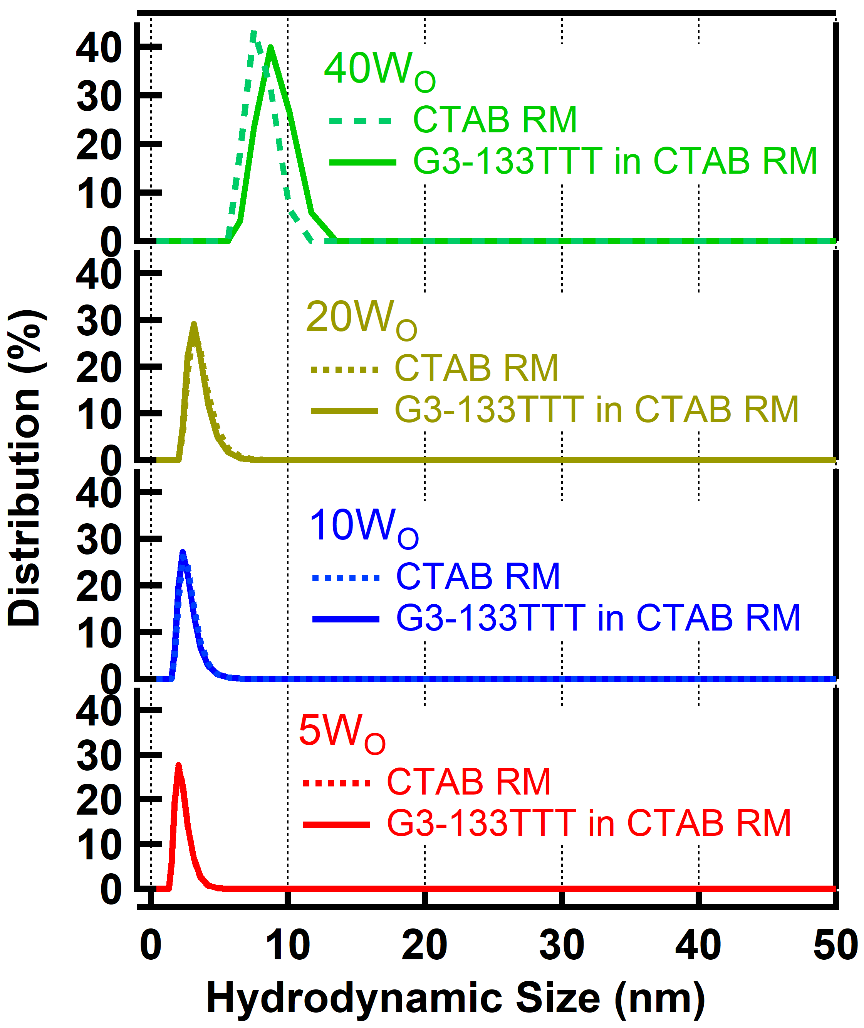

**Figure S9:** The DLS spectra of CTAB -AOT RM without and with G3-133TTT DNA (5µM) sequence.

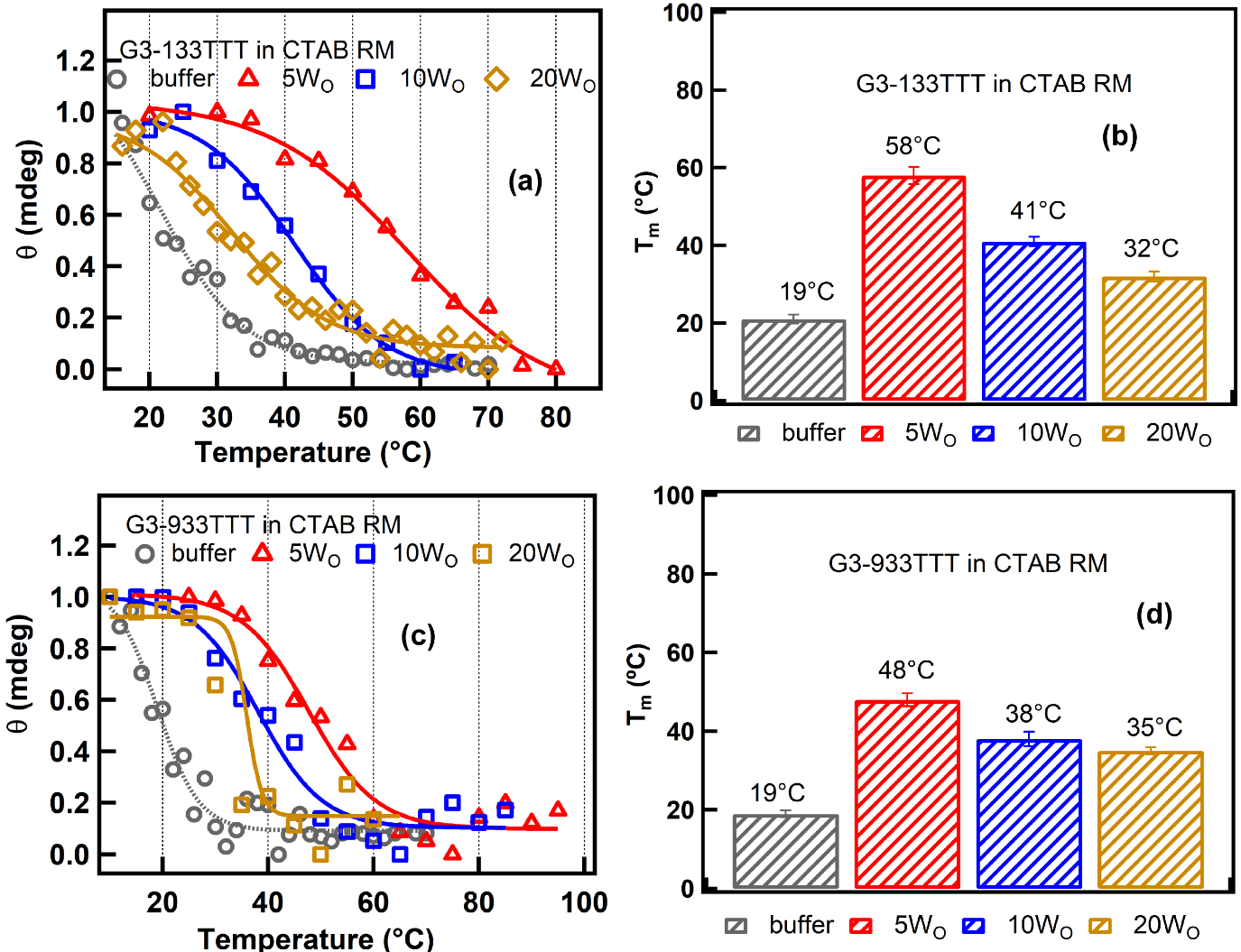

**Figure S10:** The melting curves and the melting temperature of G133TTT (a and b) and G-933TTT (c and d) in buffer and CTAB-RMs of the different water content.

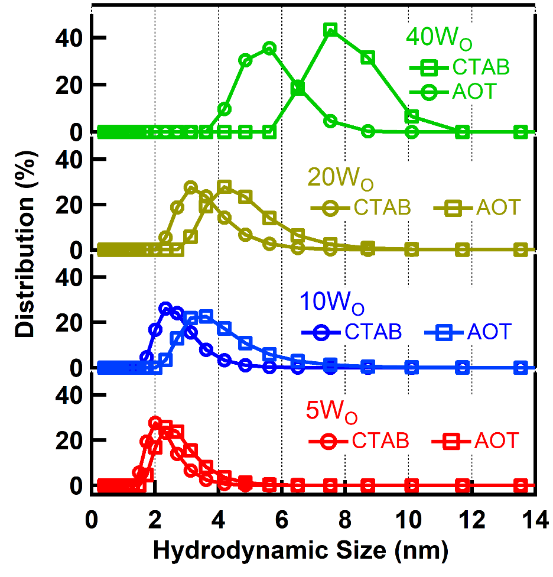

**Figure S11:** The DLS spectra of CTAB and AOT-RMs having different water contents.

**Table S1:** The amplitude, life time (ns) and the average life time (ns) of donor FAM and acceptor TAMRA attached with the G3-133TTT (a), G3-333TTT (b), G3-533TTT (c), and G3-933TTT DNA sequences in buffer and the presence of AOT-RM with different water content.

| **(a) G3-133TTT, emission of FAM @520** | | | | | | | |
| --- | --- | --- | --- | --- | --- | --- | --- |
|  | α_1_ | τ_1_ | α_2_ | τ_2_ | α_3_ | τ_3_ | τ_avg_ |
| buffer | 0.16 | 0.96 | 0.62 | 3.37 | 0.22 | 0.09 | 2.26 |
| 5Wo | 0.01 | 0.74 | 0.16 | 2.88 | 0.84 | 0.03 | 0.49 |
| 10Wo | 0.02 | 0.18 | 0.24 | 3.04 | 0.74 | 0.03 | 0.76 |
| 20Wo | 0.03 | 0.91 | 0.29 | 3.46 | 0.68 | 0.04 | 1.06 |
| 40Wo | 0.05 | 0.86 | 0.35 | 3.42 | 0.61 | 0.04 | 1.26 |
| **G3-133TTT, emission of TAMRA @580** | | | | | | | |
| buffer | 0.39 | 1.36 | 0.11 | 0.26 | 0.49 | 3.35 | 2.20 |
| 5Wo | 0.34 | 1.41 | 0.52 | 3.32 | 0.14 | 0.29 | 2.25 |
| 10Wo | 0.35 | 1.34 | 0.16 | 0.28 | 0.49 | 3.31 | 2.14 |
| 20Wo | 0.36 | 1.41 | 0.18 | 0.29 | 0.46 | 3.51 | 2.17 |
| 40Wo | 0.35 | 1.24 | 0.47 | 3.26 | 0.19 | 0.25 | 2.01 |

| **(b) G3-333TTT, emission of FAM @520** | | | | | | | |
| --- | --- | --- | --- | --- | --- | --- | --- |
|  | α_1_ | τ_1_ | α_2_ | τ_2_ | α_3_ | τ_3_ | τ_avg_ |
| buffer | 0.29 | 1.61 | 0.55 | 3.61 | 0.16 | 0.19 | 2.48 |
| 5Wo | 0.04 | 0.54 | 0.33 | 2.97 | 0.64 | 0.03 | 1.02 |
| 10Wo | 0.04 | 0.36 | 0.35 | 3.07 | 0.61 | 0.03 | 1.11 |
| 20Wo | 0.04 | 0.38 | 0.39 | 3.27 | 0.57 | 0.02 | 1.30 |
| 40Wo | 0.06 | 0.37 | 0.45 | 3.32 | 0.49 | 0.02 | 1.53 |
| **G3-333TTT, emission of TAMRA @580** | | | | | | | |
| buffer | 0.34 | 1.46 | 0.57 | 3.22 | 0.09 | 0.28 | 2.35 |
| 5Wo | 0.22 | 1.52 | 0.69 | 3.39 | 0.09 | 0.41 | 2.71 |
| 10Wo | 0.41 | 1.82 | 0.51 | 3.61 | 0.08 | 0.36 | 2.62 |
| 20Wo | 0.44 | 1.85 | 0.46 | 3.65 | 0.09 | 0.41 | 2.52 |
| 40Wo | 0.31 | 1.42 | 0.60 | 3.33 | 0.08 | 0.32 | 2.46 |

| **(c) G3-533TTT, emission of FAM @520** | | | | | | | |
| --- | --- | --- | --- | --- | --- | --- | --- |
|  | α_1_ | τ_1_ | α_2_ | τ_2_ | α_3_ | τ_3_ | τ_avg_ |
| buffer | 0.15 | 1.33 | 0.71 | 3.23 | 0.15 | 0.17 | 2.52 |
| 5Wo | 0.02 | 0.38 | 0.19 | 3.06 | 0.78 | 0.01 | 0.59 |
| 10Wo | 0.02 | 0.36 | 0.25 | 2.75 | 0.73 | 0.03 | 0.72 |
| 20Wo | 0.03 | 0.32 | 0.27 | 3.16 | 0.70 | 0.01 | 0.87 |
| 40Wo | 0.06 | 0.31 | 0.37 | 2.94 | 0.57 | 0.01 | 1.11 |
| **G3-533TTT, emission of TAMRA @580** | | | | | | | |
| buffer | 0.28 | 1.44 | 0.63 | 3.21 | 0.09 | 0.28 | 2.45 |
| 5Wo | 0.28 | 1.78 | 0.67 | 3.79 | 0.05 | 0.31 | 3.05 |
| 10Wo | 0.30 | 1.89 | 0.64 | 3.74 | 0.06 | 0.37 | 2.98 |
| 20Wo | 0.35 | 1.84 | 0.58 | 3.74 | 0.07 | 0.36 | 2.83 |
| 40Wo | 0.29 | 1.45 | 0.63 | 3.53 | 0.07 | 0.32 | 2.67 |

| **(d) G3-933TTT, emission of FAM @520** | | | | | | | |
| --- | --- | --- | --- | --- | --- | --- | --- |
|  | α_1_ | τ_1_ | α_2_ | τ_2_ | α_3_ | τ_3_ | τ_avg_ |
| buffer | 0.11 | 1.51 | 0.78 | 3.39 | 0.11 | 0.21 | 2.83 |
| 5Wo | 0.06 | 0.39 | 0.38 | 3.13 | 0.56 | 0.02 | 1.22 |
| 10Wo | 0.08 | 0.87 | 0.41 | 3.58 | 0.51 | 0.02 | 1.55 |
| 20Wo | 0.07 | 0.36 | 0.54 | 2.91 | 0.39 | 0.05 | 1.62 |
| 40Wo | 0.09 | 0.38 | 0.59 | 2.96 | 0.31 | 0.03 | 1.79 |
| **G3-933TTT, emission of TAMRA @580** | | | | | | | |
| buffer | 0.26 | 1.56 | 0.63 | 3.40 | 0.10 | 0.26 | 2.57 |
| 5Wo | 0.24 | 1.95 | 0.70 | 3.89 | 0.06 | 0.40 | 3.22 |
| 10Wo | 0.27 | 1.80 | 0.67 | 3.69 | 0.06 | 0.35 | 2.97 |
| 20Wo | 0.28 | 1.66 | 0.65 | 3.55 | 0.07 | 0.34 | 2.79 |
| 40Wo | 0.32 | 1.54 | 0.60 | 3.49 | 0.08 | 0.32 | 2.61 |

**Table S2**: The comparison of the melting temperature (T_m_ ℃ ) of the different G4 DNA sequences in the presence of the AOT and CTAB-RMs having different water content.

| **G3-133TTT** | | |
| --- | --- | --- |
| Water content | T_m_ (℃) in AOT-RM | T_m_ (℃) in CTAB-RM |
| 5Wo | >100 | 58± 2 |
| 10Wo | >100 | 41± 1 |
| 20Wo | 79± 3 | 32± 1 |
| **G3-333TTT** | | |
| 5Wo | >100 | 61± 3 |
| 10Wo | >100 | 55± 2 |
| 20Wo | 69 ± 4 | 47 ± 1 |
| **G3-933TTT** | | |
| 5Wo | 67± 2 | 48± 2 |
| 10Wo | 52± 1 | 38± 2 |
| 20Wo | 34± 1 | 35± 1 |
